# Ensemble automated approaches for producing high quality herbarium digital records

**DOI:** 10.1101/2024.02.19.580800

**Authors:** Robert. P. Guralnick, Raphael LaFrance, Julie M. Allen, Michael W. Denslow

**Affiliations:** Florida Museum of Natural History, University of Florida, Gainesville, Florida, USA; Department of Biological Sciences, VirginiaTech, Blacksburg, Virginia, USA

## Abstract

One of the slowest steps in digitizing natural history collections is converting labels associated with specimens into a digital data record usable for collections management and research. Recent work has shown a path for semi-automated approaches that can find labels, OCR them and convert the raw OCR text into digital data records. Here we address how raw OCR can be converted into a digital data record via extraction into standardized Darwin Core fields. We first showcase development of a rule-based approach and compare outcomes with a large language model-based approach, in particular ChatGPT4. We next quantified error rates in a set of OCRed labels, determining omission and commission errors for both approaches and documenting other issues. For example, we find that ChatGPT4 will often create field names that are not Darwin Core compliant. Our results suggest that these approaches each have different limitations. Therefore, we developed an ensemble approach that utilizes outputs from both in order to reduce problems from each individual method. An ensemble method reduces issues with field name heterogeneity and strongly reduces information extraction errors. This suggests that such an ensemble method is likely to have particular value for creating digital data records, even for complicated label content, given that longer labels, though more error prone, are still successfully extracted. While human validation is still much needed to ensure the best possible quality, we showcase working solutions to speed digitization of herbarium specimen labels that are likely usable more broadly for all natural history collection types.

## Introduction

The natural history collections community has made enormous progress in large scale digitization of specimens over the past two decades, catalyzed by a series of technical and social advancements (Hedrick et al., 2020). However, label digitization, a process which converts analog information on labels into digital text that can then be atomized into proper fields in digital databases, remains one of the slowest steps in overall workflows (Guralnick et al. 2024). This step has remained slow because it has required significant human input to deliver high quality results, even when collections employ some automated steps, such as label Optical Character Recognition (OCR).

Automated approaches hold promise to help speed label digitization (Takano et al., 2024). The goal of such approaches is to take an image of a label and return a high-quality output conforming to a standardized specimen record, e.g. conforming to the Darwin Core standard (Wieczorek et al., 2012; Figure 1). Unfortunately, all steps of this process often still produce relatively high error rates, such that efforts needed to correct mistakes are time costly (Guralnick et al., 2024). Adopting new automation approaches has to clear the bar of being better and faster than doing it via human effort, and dramatically shift the needle - to be much better and much faster - for there to be broad-scale uptake. Very recently, new machine learning approaches, especially Large Language Models, hold promise to dramatically improve some steps of this process, especially atomizing text into standardized fields (Weaver et al., 2023; Figure 1).

**Figure 1.**
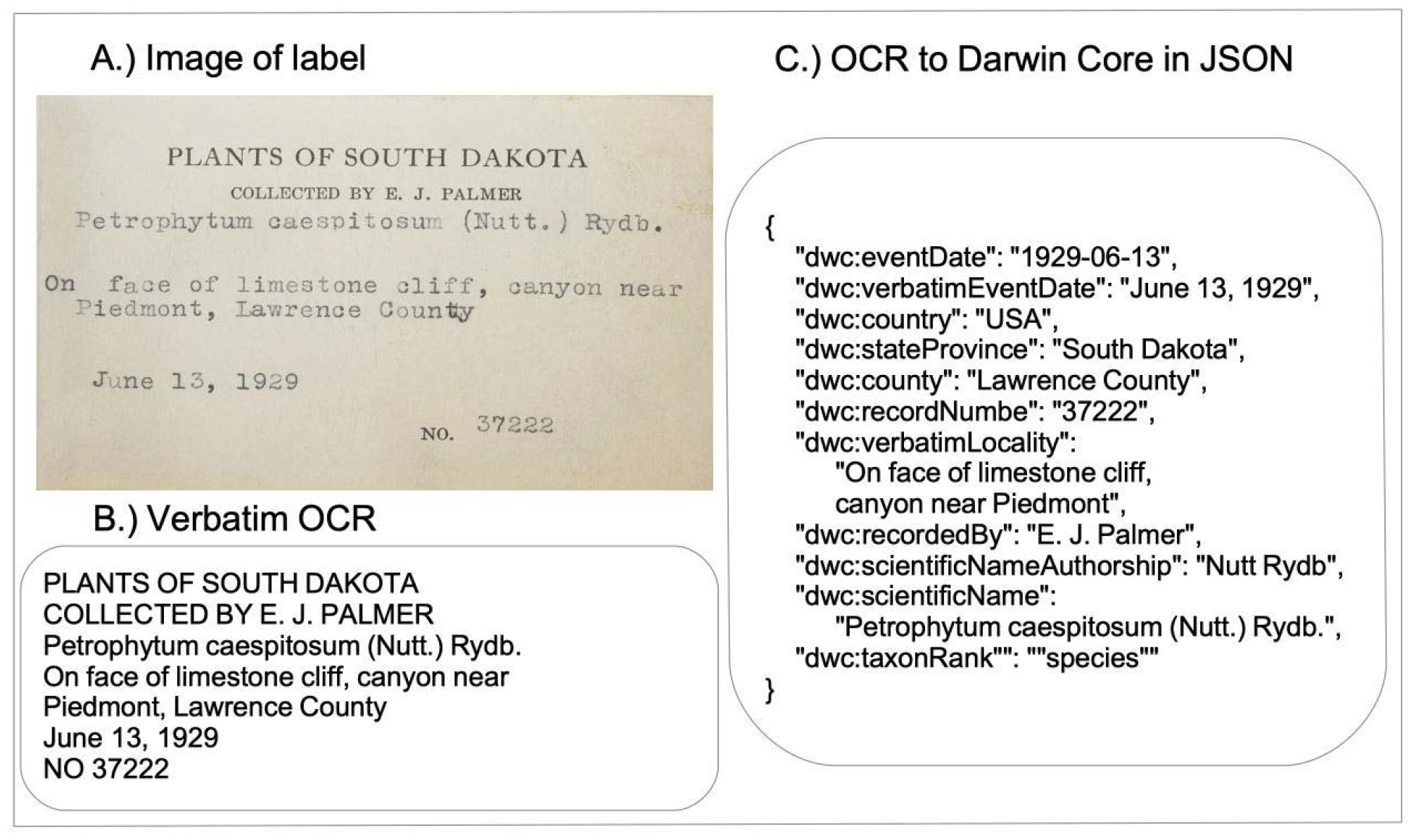
Example showing the goal of automated label digitization via the conversion of (A) label text to (B) OCR and (C) the conversion from verbatim text to JSON-formatted parsed data in Darwin Core format.

Despite their enormous potential, how much new tools such as Large Language Models (LLM), e.g. ChatGPT4 (OpenAI, 2023), can enhance the quality and speed of herbarium label digitization is just beginning to be explored (Weaver et al., 2023). Further, there are other approaches that utilize Natural Language Processing approaches (see Owen et al., 2020) that have yet to be fully tested and compared to LLM approaches. For example, rule-based natural language processing (RB-NLP) has been used in many different applications in text mining biological data and may prove to be a more reliable alternative (Xu et al., 2021). The key questions yet to be fully addressed are about the error rates in different approaches, and if and how those approaches can be combined to further improve results.

Here we provide a detailed assessment of how well different approaches work for atomizing OCR text from herbarium labels into Darwin Core fields, a standard widely used by the natural history collections community. We do so by first providing details on the development of a rule-based NLP information extraction approach and how well it performs, comparing it directly to results from finely tuned queries for extracting and atomizing label data using ChatGPT4. We calculate omission and commission error rates for both tools, focusing on core target fields that are essential to capture from labels. Finally, we showcase an ensemble approach that combines rule-based and ChatGPT outputs, which performed far better than either approach separately. Our overall work provides an assessment of what is possible, keeping in mind we are just at the start of what is likely to be a major transition from human transcription to more efficient automated approaches in label digitization efforts.

## Material and Methods

### OCR test data

We used a set of OCR’ed label data from Guralnick et al. (2024) as a test set. All of the labels came from Global Biodiversity Information Facility (GBIF). We searched the GBIF database in September 2023 for all specimens meeting the following criteria; members of Tracheophyta, collected in the United States, record containing a specimen image, preserved specimen type, english language and not cultivated. This search resulted in 4,091,778 records of which we randomly selected 2,128, most of which were from the main label on the specimen sheet. These labels were OCR’ed using a custom pipeline that has a set of pre- and post-processing steps that improves quality over off the shelf Open-Source solutions such as Tesseract (https://github.com/tesseract-ocr/tesseract). The OCR content was not corrected prior to being used in downstream workflows, because the goal was to determine success of parsing when there are an unknown number of OCR errors in the input.

### Rule-based NLP development

Our label rule-based NLP extraction approach (abbreviated LRBE) uses a multistep approach to extract text and link to named Darwin Core terms. The rules themselves are written using spaCy (Honnibal and Montani, 2017), which is a key Natural Language Processing library in the Python programming language, with enhancements we developed to streamline the rule building process. The general outline of the rule building process follows:

1. We assembled Darwin Core terms names that need to be associated with parts of OCR content. We then used existing term labels already formatted in Darwin Core format from iDigBio and other sources with a set of content terms associated with those labels.
2. We also assembled some key corpi e.g. names of known plant taxa names from Kew Plants of the World online, the WFO plant list, and the ITIS database to help with matching to key fields, e.g. dwc: scientificName.
3. We used all this expert content to develop spaCy’s phrase matchers. These are rarely sufficient for capturing the content needed, especially for more complex content such as locality or habitat, so we used phrase matches as anchors for extracting more complex content.

3. We then built more complex content from the simpler matched phrases using spaCy’s rule-based matchers repeatedly (Figure 2). For related traits, we then linked the traits to each other (entity relationships) using spaCy rules.

In developing the LRBE we iteratively tested the performance of the tool on the same set of OCR labels. This involved RP and MD examining 100s of already formatted labels and finding common problems with extraction quality, and then determining whether there was a rule that could be added to improve the results. After multiple iterations, we were able to remove many potential issues and improve parsing, but the challenge remained that the LRBE often produced extracted content with both commission and omission errors. We quantified those errors and compared them with rates from ChatGPT4 and an ensemble approach below.

**Figure 2.**
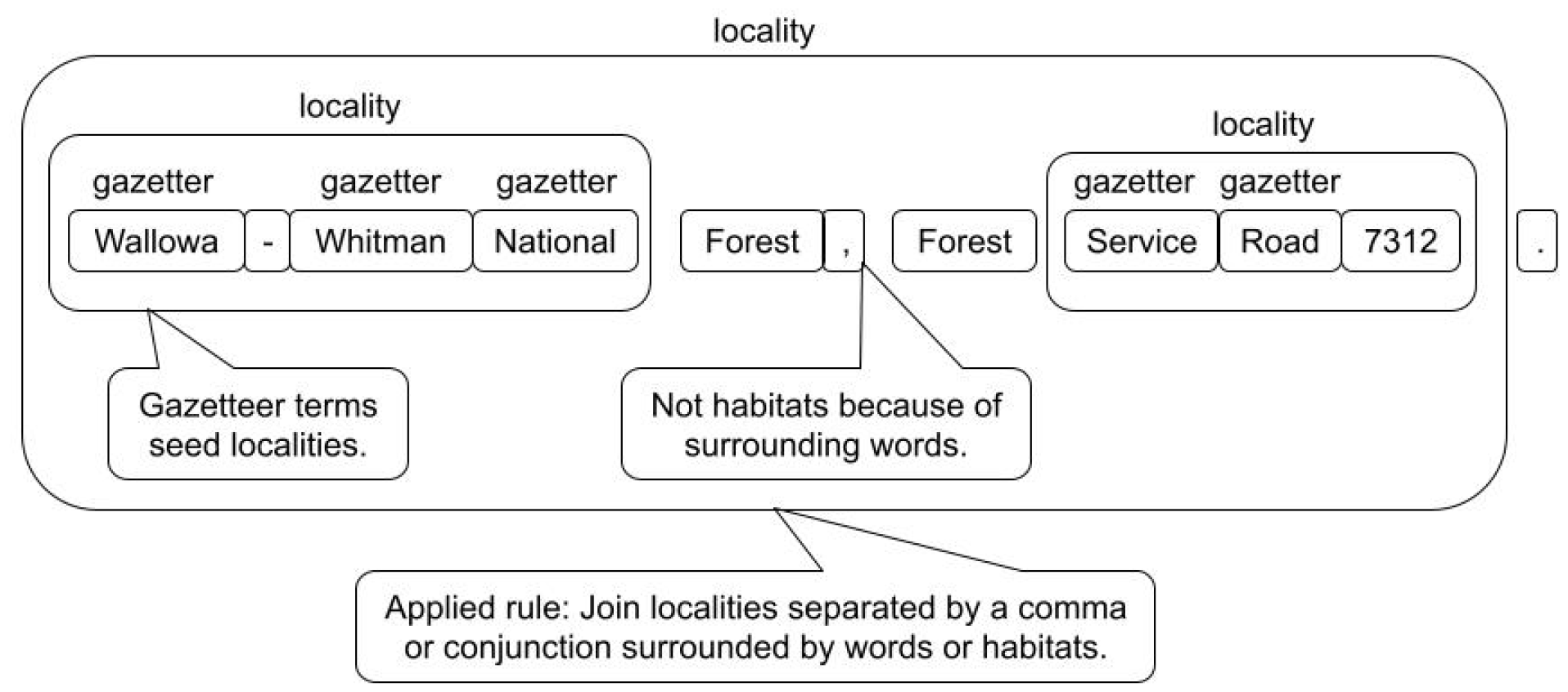
Example of a rule-based parsing approach to discover locality descriptions within a label. The RLBE starts by using a Gazetteer to find key works that seeds the locality. A set of rules are used to extract all the content that belongs in the field “dwc:locality”. For example, in this case ‘Forest’ is not a habitat term in either usage above, because in the first case it is tied to the “National” that precedes it, which is a place name and in the second post-ceded by “Service” which implies a type of road. Different locality pieces are joined based on an applied rule to create a final output string labelled as “locality”.

### Using ChatGPT for label parsing

In contrast to the LRBE, large language models like ChatGPT require very little knowledge of how they work and most of the upfront effort is with “prompt engineering”. Prompt engineering is shaping your queries to the large language model so that they yield the best results possible; an art form in itself. Our approach to prompt engineering was to keep the prompts small and focused on extracting information in Darwin Core format.

The prompt that we used for this paper was “Extract all information from the herbarium label text and put the output into JSON format using DarwinCore fields including dynamicProperties” followed by the same OCR label text used for testing the LRBE. This is a small prompt that worked reasonably well.

### Developing an Ensemble approach

ChatGPT4 and the LRBE had their own strengths and weaknesses. Because of this, we opted to ensemble the two approaches with the goal of utilizing the best of both approaches to reduce error rates. Doing so involved a process of reconciling outputs across the same or similar fields to a consensus output. We refer to the overall process as ensembling, but within a field or set of fields, we call this process reconciliation. With some exceptions, there is one reconciler per Darwin Core term. Most reconcilers are very simple, prioritizing either ChatGPT or LRBE outputs and making sure that those align with known Darwin Core terms. Others are a bit more complex as we discuss below. Each reconciler takes as input, the JSON data from LRBE and from ChatGPT that was subsequently cleaned, and the original label text that was fed to both.

One of the most challenging problems with ChatGPT4 outputs is that it attempts to extract data into Darwin Core terms as best it can, but will often create field names for content it cannot fit in Darwin Core, some of which could be mapped to known DwC terms. This led to a surprising profusion of terms that are not in the darwin core controlled vocabulary, discussed more below in Results. To handle this data we created aliases for all the reconcilers that include non-Darwin Core terms that map them to the correct term. For example, the ChatGPT created the following terms: “dwc:locationState”, “dwc:state”, and “dwc:province” that we aliased to the proper term “dwc:stateProvince”.

An example of a challenging Darwin Core field to reconcile is “dwc:verbatimLocality”. Before developing the reconciler for this field we noted a few observations: 1) ChatGPT tends to correctly find the locality more often than LRBE but when LRBE finds the correct locality, it often finds a longer correct version. This longer version is often broken up into a list of locality phrases rather than one contiguous locality value; 2) ChatGPT sometimes puts a separate locality notation under the “dwc:locationRemarks” term; 3) ChatGPT’s version of locality is sometimes presented as a nested object. That is, it is itself a dictionary of locality related terms that need to be assessed and reconciled.

Given the above, the process we use to reconcile “dwc:verbatimLocality” is as follows. First, we look for the locality in the ChatGPT output listed under any of its aliases and in dwc:locationRemarks. If ChatGPT’s version of locality is a nested object then try to pull a good locality from one of its sub-terms. If that was not possible we then use the LRBE’s version. When examining LRBE’s version, we focus on DwC locality fields because LRBE does not have the same issues as ChatGPT with inventing DwC fields. If the LRBE content is a contiguous list of partial localities, we composite that list into a seamless single string. In cases where ChatGPT locality (or aliases) are present, we still check in the LRBE version and if the ChatGPT is completely contained within a larger LRBE locality string, we then use the LRBE version. Finally, we check to see if content in location remarks from ChatGPT can be used to either extend the currently used locality or use it as another item in a locality list.

### Testing error rates for LRBE, ChatGPT4 and Ensemble Approaches

We randomly selected 100 total outputs from LRBE and ChatGPT and MWD and RPG, skipping labels that weren’t the main labels (see below), and scored error rates as follows. We defined a set of core fields that are often of particular importance to properly capture and are present on a majority of labels. These fields are: recordedBy, recordNumber, eventDate, locality, country, stateProvince, county, and scientificName. We explicitly captured information on the number of commission and omission errors in target fields, in order to determine performance of the text extraction tools. A commission error is a case where there is extra content in a field that is clearly not correct and belongs in another field. Omission errors are cases where the extraction approaches missed content that was clearly supposed to be in the field in question. Further, we noted many of the errors, but did not explicitly count omission and commission, for other ancillary, non-target fields. We also noted cases where ChatGPT made semantic interpretations, such as expanding a country name from “USA” to “United States of America”, but did not score those as errors. We did not flag cases where either tool returned dates in standard formats from whatever format was used on the typewritten labels. We explicitly skipped determination labels and we also skipped labels where OCR results were so poor as to significantly impact ability of either LRBE or ChatGPT4 to work effectively. We did, however, not clean OCR outputs, because the goal here is to see how well these approaches perform in automated pipelines where there is likely to be a low percentage of OCR issues still left during the parsing step.

We also scored the quality of ensembled outputs focusing on the same 100 labels already scored for the other approaches. Here MWD and RPG scored those results focusing on an overall omission and commission error rates on core fields and ancillary ones, rather than only focusing scoring on core target fields. After this first pass on all fields, we went back through and counted errors per core field individually for each field, in order to assess which core fields were most problematic. We then tabulated overall error rates across all 100 example labels for LRBE ChatGPT4 and ensembles. Finally, we addressed if length of label explains the number of total errors, which we expect since more content should mean more chances for either approach to mis-assign content and thus lead to errors in reconciled outputs. We simply fit a single predictor model with length of the label, measured as total number of characters, as a predictor of total error rate, using the lm() function in base R (R Core Team, 2021).

### Testing ChatGPT4 Darwin Core field names

ChatGPT does not always return canonical Darwin Core field names published as part of the standard (https://dwc.tdwg.org/terms/). In order to quantify the magnitude of this problem, we counted how many times ChatGPT used a non-canonical field name from our full record set. We counted in particular how many unique field names that ChatGPT “invented” that were non-canonical and the number of records and fields that were impacted across all the labels. Finally, for the core fields defined above, we determined how many synonyms existed that linked to the proper core field name.

## Results

### Summary of performance within core fields

Table 1 summarizes the commission and omission error rates for the LRBE, ChatGPT4 and ensemble approaches for core fields defined above in the Methods. The key finding is that ChatGPT performs much better than rule-based parsing in terms of omission and commission errors, with over 2-fold fewer errors than a rule-based approach. ChatGPT4 is particularly good in not making commission errors, with a nearly three-fold improvement in commission. In the case of the LRBE, multiple dates and especially numbers (such as road numbers in locality descriptions) often ended up wrongly linked to eventDate or recordNumber leading to a much higher rate of commission. Ensemble methods performed the best of all, significantly reducing omissions and commission errors compared to the ChatGPT4 or LRBE. The drop in omission and commission rates and improvement in reducing non-Darwin Core terms is due to complementarity and leveraging what both tools do best. LRBE is less likely to hallucinate terms while ChatGPT is better at extraction, but one or the other will often do better on different labels. The ensemble often captures the best outcomes. An example output from both ChatGPT4, LRBE and ensembles, is shown in Figure 3. That example nicely illustrates issues with ChatGPT4 including omission errors and Darwin Core field names issues and commission issues with LRBE, all of which are resolved in the ensemble output. We discuss more regarding the performance of both approaches and ensembles below.

**Table 1.** Error counts for ChatGPT4, LRBE and ensembled results based on scoring of 100 randomly selected herbarium records. These are errors for essential Darwin Core fields when present (recordedBy, recordNumber, eventDate, locality, scientificName, country, stateProvince, county). For the randomly selected 100 records used there are 723 possible errors since not all fields are present in each record.

| Method | ChatGPT4 | Label Rule-based | Ensemble-based |
| --- | --- | --- | --- |
| Commission Errors | 23 | 83 | 12 |
| Omission Errors | 43 | 64 | 32 |
| Total | 66 | 147 | 44 |

**Figure 3.**
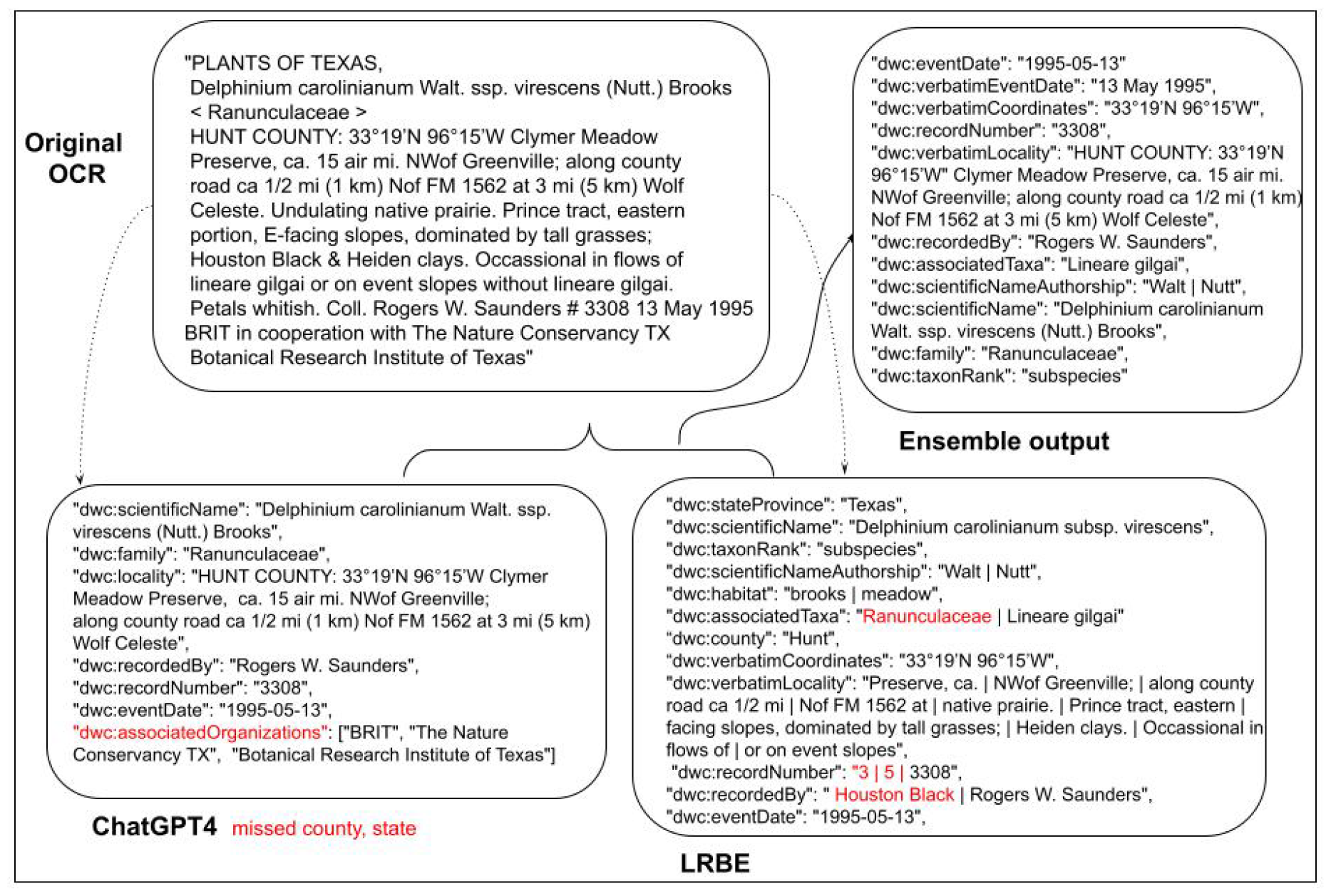
Examples of the workflow and processing outputs from an exemplar label, starting with the original OCR shown top left, the ChatGPT4 output (bottom left), the LRBE output (bottom right) and ensemble output (top right). We have excised any dynamicProperties content returned by either extraction method. Errors in extraction are shown in red, with omissions shown below label contents. In the ChatGPT4 output, dwc:associatedOrganizations is an example of a hallucination as this is not a Darwin Core term.

We also calculated overall error rates for the ensemble approach, ignoring more free-form content that doesn’t fit into typical Darwin Core fields e.g. flower color, which is often captured in the wastebasket field “dwc:dynamicProperties”. The ensemble approach performs well on these ancillary fields as well, with only 11 errors recorded across all of these fields for our 100 label test set. Because we have a full set of errors (excluding content placed in dynamicProperties), we were able to run a simple test to determine if longer labels (measured as number of characters) are more likely to have more errors. Our simple linear model using label length as a predictor of overall error rate is significant (p<.001) but the overall adjusted R^2^ is .12, suggesting only a modest relationship between label length and error rate.

Finally, we also examined ensemble error rates across core fields (Table 2). These results surprisingly show that the most errors in the ensemble approach are in eventDate and stateProvince. While locality also had a relatively large number of errors, the ensemble-based method is surprisingly good at assembling coherent locality information from these records, which a priori was anticipated to be a significant challenge since localities are often complex sets of information that need to be assembled from the label text. We checked specific instances of eventDate omission, and the key issue with both LRBE and ChatGPT4 was not recognizing certain event date formats, especially of the form “9-IV-1977”, with roman numerals for months. For stateProvince and county, the most common error - an omission - was state, province or county being included in locality and not pulled out into its own field.

**Table 2.**
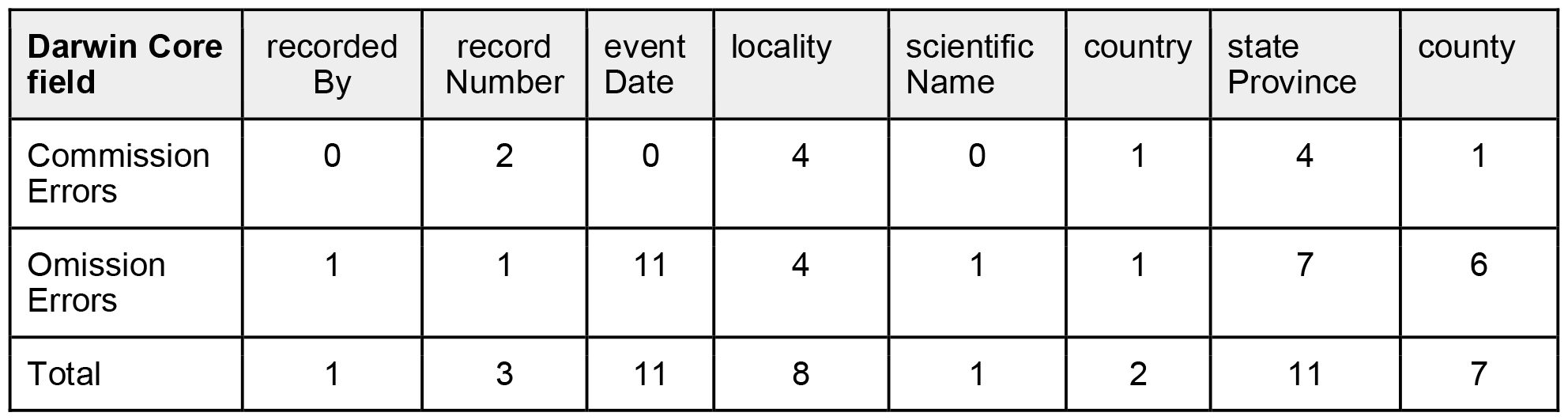
Types of core field errors contained in reconciled output. Not all core fields are present in each record. 44 total errors in 100 assessed records. We use Darwin Core camel case names for these field names.

| Darwin Core field | recorded By | record Number | event Date | locality | scientific Name | country | state Province | county |
| --- | --- | --- | --- | --- | --- | --- | --- | --- |
| Commission Errors | 0 | 2 | 0 | 4 | 0 | 1 | 4 | 1 |
| Omission Errors | 1 | 1 | 11 | 4 | 1 | 1 | 7 | 6 |
| Total | 1 | 3 | 11 | 8 | 1 | 2 | 11 | 7 |

### ChatGPT4 and Darwin Core field names

Out of 2128 herbarium labels fed to ChatGPT, it extracted 420 terms, of which 155 were valid Darwin Core terms, and 265 were hallucinated. Note that this is after performing a term cleanup pass on the data and ignoring any changes to the case of the letters. Although ChatGPT hallucinated a large number of terms, the actual number of instances of fields across labels that used the hallucinated terms was not as large. There were 23,094 instances extracted, but the number of those that had hallucinated term labels was only 1,062 (or 4.6%). Hallucinations were unpredictable. Some assumed there was another namespace that existed e.g. “gbif:identificationRemarks”, although dwc:identificationRemarks is a valid term. Other hallucinations were more esoteric, such as “QF”. In cases such as “gbif:identificationRemarks”, the term name can be converted to the correct one and used. In the case of “QF”, there is little to be done besides attempting to examine the content associated with that term.

One key concern with ChatGPT4 apart from the hallucinated terms is the formatting of the return. The returned JSON output was often improperly formatted with things like extra commas, improper quoting, the replacement or addition of extra characters, and adding extra text surrounding the JSON output. This problem was more common. Of the 2128 labels that were processed 503 (or 23.64%) of them had data formatting issues. The end result is that 503 (or 23.64%) have bad JSON that we could salvage, 542 (or 25.47%) labels have one or more hallucinated terms, and there is an overlap of 112 (5.26%) labels that have both.

## Discussion

This work showcases the power of using multiple different approaches for producing digital data records that together help reduce errors that are found using any single method. In particular, we developed a rule-based NLP approach (LRBE), and tested how well it performed against a well-used large language model, ChatGPT4. The LRBE approach can have excellent precision, and sometimes excellent accuracy and critically it will neither hallucinate names of fields or instance value data in those fields, and it always produces the exact same result on repeated uses. It is also easily adjusted when improvements are needed. However, the LRBE also requires significant effort by an expert that understands both herbarium label construction and natural language processing tasks well enough to correct issues. The need for this significant effort is due to having to write one or often several parsing rules for every field type and form. For instance, when extracting information about taxa there are patterns for every commonly used taxon level, and separate functions for when there is a binomial or trinomial term versus a monomial term. The taxon authority extraction builds on the bi-, tri-, or monomial term. This is then fed into a function that recognizes a binomial taxon followed by an authority which is then followed by a lower-level term with its own authority, like, “Neptunia gracilis Muhl. ex Willd. var. varia (Nutt.) Brewer”. There are even other forms for taxon names such as “Neptunia gracilis & Mimosa sensitiva” or when one species is mentioned in relation to another like, “It resembles M. sensitiva in amplitude”. The end result is that development of a working LRBE is both time intensive and challenging. By contrast, ChatGPT4 requires a relatively simple set of prompt engineering approaches and can produce digital data records in Darwin Core format that contain fewer errors. However, ChatGPT4 has other issues, e.g. non-deterministic results and the potential for a profusion of terms that do not conform to existing standards.

Ensemble approaches using outputs from rule-based NLP and from ChatGPT4 can help resolve both issues, as shown in our results. The ensemble approach leverages strengths of the RLBE in terms of mostly assembling the right content into a predefined set of known Darwin Core fields. This provides a useful scaffold and framework to blend in the improved results from ChatGPT4. While it is possible that far more careful prompt engineering could also improve ChatGPT4 results, it is challenging to understand *a priori* what prompts are required. Given that ChatGPT4 has multiple issues, ensembling fixes many of them simultaneously, and requires little extra overhead, we advocate the value of this multimethod approach as a pragmatic step forward. We should note that ensembling doesn’t obviate the need for data consistency checks or formatting corrections from ChatGPT4. This means that simply using out of the box large language models for automated assembly of digital data records is likely to be far more challenging than first anticipated, even if there are collections staff with informatics expertise.

We note that our work here focuses mostly on core fields that are often required for capture for best use of collections downstream for collections management purposes and for research. Herbarium labels often also report key traits of the specimen, such as flower color or leaf size. Here as well it can be challenging to capture traits successfully, even though LRBE has its origins in trait parsing. Part of the issue is that herbarium labels are often rich with information that can ultimately confuse a LRBE approach. The longer the label - the more content it reports - the easier it is for extraction tools to pull the wrong content and associate it with the wrong term, although this is one of many factors that likely impacts how well information extraction works. For instance, LRBE will sometimes mistake route numbers like “Rt. 12” for a count. This can be counteracted by adding more rules that bar a count when it is preceded by a route abbreviation. Trying to build rules for all possible vagaries of how labels are written is an impossible task and the key goal is to find common errors of commission and reduce their rate as much as possible using such rules.

We close here with four key observations about the current state of automated label digitization and likely next steps. First, ChatGPT4 is a commercial solution and costs money to use. A longer term solution will be developing and deploying open source large language models such as LLaMA (Touvron et al., 2023). Such models can be tuned to perform significantly better than a generalized LLM such as ChatGPT4. Second, automated label parsing cannot “solve” the problem of label digitization. While continuing advances in quality of OCR of labels and improving data extraction will likely further reduce error rates (as shown for OCR in Guralnick et al., 2024), and we believe these rates can potentially get close to the quality of human effort, there will always be the need for humans to validate and improve the quality of such extraction. We have made the case in other work (Guralnick et al., 2024) of the importance of humans in the loop both for quality improvement and to improve model performance and we advocate strongly for that approach here. Finally, we believe the work here can be extended and utilized broadly for natural history collections digitization. One aspect of this extension is recognizing that automated approaches can be combined to deliver both digital data records and other insights, such as leaf traits, from specimens simultaneously (Weaver and Smith, 2023). Our work focused on OCR of typewritten labels (Guralnick et al., 2024). Future advancements could focus on the integration of handwritten text recognition (HTR) in an attempt to improve information extraction for older labels which were generated prior to the use of typewriters and computers. More broadly, herbarium labels are some of the most verbose, commonly containing heterogeneous content, when compared across different types of natural history collections. By contrast, insect labels are typically far less verbose or heterogeneous, and therefore likely less error prone, for automated extraction approaches. Further efforts to test approaches across label types and building more production strength tools are a critical next step.

## Data and code availability

All code is open source and available at https://github.com/rafelafrance/digi-leap Data and scoring sheets used in creating and assessing models are available at https://zenodo.org/uploads/10642072 [draft, not yet public]

## Acknowledgements

This work was supported by a grant to RPG and JMA from the National Science Foundation #2027234 and #2027241 entitled “Leaping the Specimen Digitization Gap: Connecting Novel Tools, Machine Learning and Public Participation to Label Digitization Efforts”. As part of this larger funded effort, we have received extensive data entry, contributions to training datasets and validation from many conscientious and dedicated Notes from Nature volunteers, for whom we are extremely grateful. We appreciate the help of the steering committee who has helped advise on this grant: Jason Best, Libby Ellwood, Sharon Grant, Deborah Paul, Melissa Tulig, and Jenn Yost. We also acknowledge our other key collaborators on this work: Samantha Blickhan, Nico Franz, Edward Gilbert, Nelson Rios, and John Wieczorek.

## Literature Cited

Hedrick, B. P., Heberling, J. M., Meineke, E. K., Turner, K. G., Grassa, C. J., Park, D. S., Kennedy, J., Clarke, J. A., Cook, J. A., Blackburn, D. C., Edwards, S. V., & Davis, C. C. (2020). Digitization and the Future of Natural History Collections. Bioscience, 70(3), 243–251.

Honnibal, M., & Montani, I. (2017). spaCy 2: Natural language understanding with Bloom embeddings, convolutional neural networks and incremental parsing.

Guralnick, R., LaFrance, R., Denslow, M., Blickhan, S., Bouslog, M., Miller, S., Yost, J., Best, J., Paul, D. L., Ellwood, E., Gilbert, E., & Allen, J. (2024). Humans in the loop: Community science and machine learning synergies for overcoming herbarium digitization bottlenecks. Applications in Plant Sciences. 10.1002/aps3.11560.

OpenAI: Achiam, J., Adler, S., Agarwal, S., Ahmad, L., Akkaya, I., Aleman, F. L., Almeida, D., Altenschmidt, J., Altman, S., Anadkat, S., Avila, R., Babuschkin, I., Balaji, S., Balcom, V., Baltescu, P., Bao, H., Bavarian, M., … Zoph, B. (2023). GPT-4 Technical Report. In arXiv [cs.CL]. arXiv. http://arxiv.org/abs/2303.08774

Owen, D., Livermore, L., Groom, Q., Hardisty, A., Leegwater, T., van Walsum, M., Wijkamp, N., & Spasić, I. (2020). Towards a scientific workflow featuring Natural Language Processing for the digitisation of natural history collections. Research Ideas and Outcomes 6:e55789.

R Core Team (2021). R: A language and environment for statistical computing. R Foundation for Statistical Computing, Vienna, Austria. URL https://www.R-project.org/.

Takano, A., Cole, T. C. H., & Konagai, H. (2024). A novel automated label data extraction and data base generation system from herbarium specimen images using OCR and NER. Scientific Reports, 14(1), 112

Touvron, H., Lavril, T., Izacard, G., Martinet, X., Lachaux, M.-A., Lacroix, T., Rozière, B., Goyal, N., Hambro, E., Azhar, F., Rodriguez, A., Joulin, A., Grave, E., & Lample, G. (2023). LLaMA: Open and Efficient Foundation Language Models. In arXiv [cs.CL]. arXiv. http://arxiv.org/abs/2302.13971

Weaver, W. N., Ruhfel, B. R., Lough, K. J., & Smith, S. A. (2023). Herbarium specimen label transcription reimagined with large language models: Capabilities, productivity, and risks. American Journal of Botany, 110(12), e16256.

Wieczorek, J., Bloom, D., Guralnick, R., Blum, S., Döring, M., Giovanni, R., Robertson, T., & Vieglais, D. (2012). Darwin Core: An Evolving Community-Developed Biodiversity Data Standard. PloS One, 7(1), e29715.

